# IMMClock reveals immune aging and T cell function at single-cell resolution

**DOI:** 10.1101/2024.11.13.623449

**Authors:** Yael Gurevich Schmidt, Di Wu, Sanna Madan, Sanju Sinha, Sahil Sahni, Vishaka Gopalan, Binbin Wang, Saugato Rahman Dhruba, Alejandro A. Schäffer, Nan-ping Weng, Nicholas P. Restifo, Kun Wang, Eytan Ruppin

## Abstract

The aging of the immune system substantially impacts individual immune responses, yet accurately quantifying immune age remains a complex challenge. Here we developed **IMMClock**, a novel immune aging clock that uses gene expression data to predict the biological age of individual CD8⁺ T cells, CD4⁺ T cells, and NK cells. The accuracy of IMMClock is first validated across multiple independent datasets, demonstrating its robustness. Second, utilizing the IMMClock, we find that intrinsic cellular aging processes are more strongly altered during immune aging than differentiation processes. Thirdly, our analysis confirms the strong associations between immune aging and established processes such as cellular senescence, exhaustion, and telomere length at the single cell level. Furthermore, immune aging is accelerated under several disease conditions such as type 2 diabetes, heart disease, and cancer. Finally, we apply IMMClock to analyze a perturb-seq gene activation screen of T cell functionality. We find that the post-perturbation immune age of individual T cells is strongly correlated with their pre-perturbation immune age. Furthermore, the immune age at resting state of individual T cells is strongly predictive of their post-stimulation activation state. Overall, IMMClock advances our understanding of immune aging by providing precise, single-cell level age estimations. Its future applications hold promise for identifying interventions that concomitantly rejuvenate and activate T cells, potentially enhancing efforts to counteract age-related immune decline.

## Introduction

Aging impacts organisms at multiple levels, leading to functional decline and increased susceptibility to diseases. Robust quantification of biological age is essential for understanding the underlying mechanisms of aging and for developing effective interventions^1^. Aging clocks, computational models that estimate biological age from epigenomic and transcriptomic features, such as DNA methylation and gene expression, have become valuable tools for assessing how an organism’s biological age compares to its chronological age. DNA methylation clocks^2–5^ have pioneered our efforts to build and test such aging clocks. They have been followed by transcriptomic aging clocks, which leverage gene expression patterns^6–10^, which offer another dimension for biological age estimation. Given the critical role of the immune system in maintaining overall health and its substantial alterations with age, several immune-focused aging clocks have been developed to specifically assess aspects of immune system aging^11–13^. These specialized clocks have provided important new insights into immune health and its association with disease, enhancing our ability to better understand and in the future, possibly treat age-related immune decline.

Within the immune system, T cells play a pivotal role in immune responses against infections and tumors and importantly, in modulating the response to cancer immunotherapy^14,15^. However, their capacity to proliferate and function effectively declines with age, associated with increased susceptibility to infectious diseases, reduced effectiveness of vaccines, and a diminished ability to control tumor growth^16–20^. Here we aim to advance our understanding of how aging affects T cell functionality by developing an aging clock designed to capture the heterogeneity of T cell aging at the single-cell level. Although DNA methylation clocks are highly accurate^2,3^, whole-genome methylation profiling at the single-cell level remains technically challenging, resulting in limited and sparse data for aging studies^21,22^. In contrast, single-cell transcriptomic approaches offer such an opportunity and include many validation datasets, also enabling to further study their functional implications in depth^23^. Previous work, such as studies by Buckley et al.^24^ on neurons in mice, Zhu et al.^25^ for supercentenarians, and Lu et al.^26^ on CD8^+^ T cells in humans, have already made valuable first steps in developing single-cell clocks, though these studies were performed on relatively small dataset sizes. This highlights the need to develop and study single-cell aging clocks for different immune cell types that are trained and validated on large and independent datasets, for the first time.

To this end, we developed IMMClock (**IMM**une **C**ell c**lock**), leveraging a large-scale human blood dataset that is two orders of magnitude larger than those used in previous studies. The resulting IMMClock is first comprehensively tested and validated across both single-cell and bulk expression large-scale datasets. Second, we demonstrate its utility in quantifying and validating key associations between immune aging and markers of senescence, exhaustion, and telomere length, as well as various clinical phenotypes. Most importantly, we use IMMClock to study our primary research question: how does aging influence T cell functionality? To this end, we examined the association between immune aging and T cell activation by analyzing a functional screen of genetic perturbations in T cells. Quite remarkably and surprisingly, we find that the immune ages of cells, as predicted by IMMClock, are strongly negatively correlated with the levels of T cell activation observed in these screens. IMMClock was developed using human aging data spanning many decades, while these functional screens measure changes over just a few days. Notably, we find that the immune age of resting T cells is strongly associated with their functional activation post-stimulation, offering new avenues for facilitating the development of targeted interventions to enhance T cell functionality.

## Results

### Overview of IMMClock and its performance

To build the IMMClock, we analyzed the OneK1K single-cell transcriptomics cohort^27^, which consists of approximately 1.3 million peripheral blood mononuclear cells (PBMCs) from 982 healthy donors, ranging in age from 19 to 97 years. These cells were annotated by the original authors as belonging to various immune cell types, including five major categories: CD8^+^ T cells, CD4^+^ T cells, NK cells, B cells, and monocytes. For our analysis, we focused only on these five types (**Fig. 1A, B**). Building upon this dataset, we developed IMMClock (**IMM**une **C**ell c**lock**), a machine learning model that employs an elastic net model (see **Methods**) to infer immune cell-type-specific biological ages based on single-cell transcriptomics data. We define the immune cell-type-specific biological age as “*immune age*”, which can pertain to either an entire individual or a specific single immune cell.

**Figure 1.**
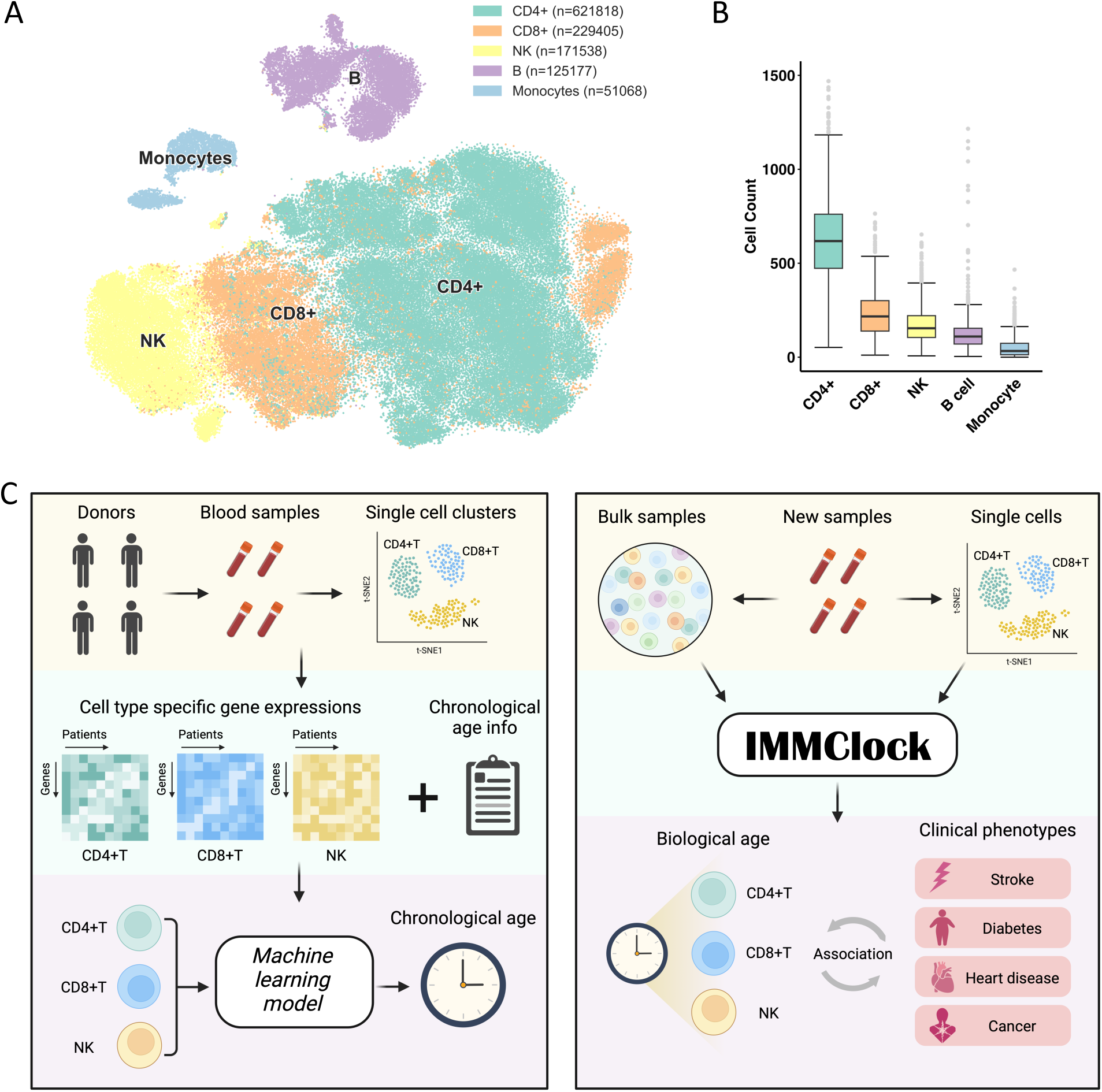
Overview of the OneK1K dataset and IMMClock. **(A)** TSNE visualization of the OneK1K single-cell dataset, highlighting the clustering of the five main immune cell types. **(B)** Bar plot showing the number of cells from each donor across the five primary immune cell types. **(C)** Diagram outlining the study design, from data collection and preprocessing to the development and application of IMMClock for immune age assessment. The left subpanel illustrates the training process for cell-type-specific IMMClocks. The right subpanel shows the testing pipeline applied to independent datasets and potential applications.

IMMClock takes the gene expression values for one individual immune cell or the average gene expression values of a particular immune cell type from a specific person as input, outputting the predicted immune age for the cell or the person, respectively. To account for cell type specificity, we developed an IMMClock independently for each cell type (**Fig. 1C**). For validation, we used data from seven other cohorts of healthy donors’ blood immune cells (**Supp. Table 1**), including single-cell RNA-seq studies such as the Asian Immune Diversity Atlas ^28^(500 individuals), two recent large datasets (Terekhova et. al, 2023^29^, 166 individuals; Perez et. al, 2022^30^, 99 individuals), and the Framingham Heart Study^31,32^, which provides bulk RNA-seq from whole blood but was chosen as it includes rich clinical phenotypic data from 2,691 healthy individuals. We first specifically assess whether IMMClock can accurately infer immune age for persons or cells, and next, whether these predictions correlate with chronological age and are relevant to specific aging-related biological processes and clinical phenotypes.

Aging in the immune system is characterized by two key processes: the differentiation of immune cells, in which naïve cells differentiate into more specialized functional types, and intrinsic aging-related changes within the same cell state/types^33,34^. Notably, the differentiation process leads to a shift from a higher proportion of naïve T cells to effector and memory T cells^35^, resulting in changes in immune cell composition, as reported by numerous studies^35–37^. The intrinsic aging process is driven by specific transcription alterations relevant to senescence within the same cell states/types^34^. We applied multiple IMMClocks to further disentangle the effects of these processes on immune system aging by conducting a comparative investigation of models trained on different cell types/states, studying their relative strength of associations with chronological age (**Fig. 2A**).

**Figure 2.**
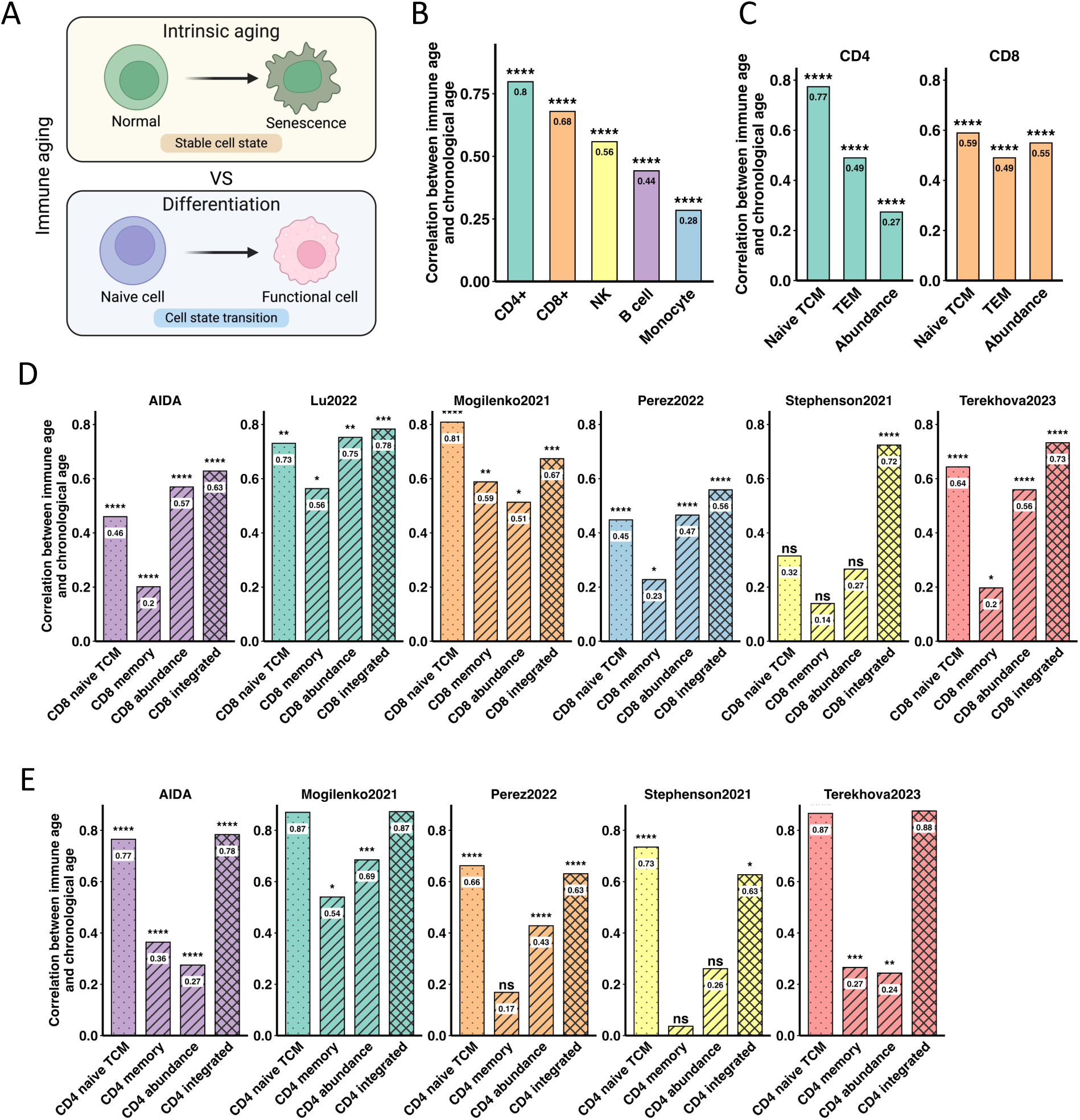
IMMClock shows high performance in predicting chronological age and enhances our understanding of immune system aging. **(A)** illustration of the two aging-related processes: the intrinsic aging process within the same cell state/type and the differentiation process that leads to compositional shifts. **(B)** Bar plot demonstrating the cross-validation performance of IMMClock using the integrated model on the OneK1K dataset. The Y-axis represents the Spearman correlation between the predicted immune age and chronological age at the person level. **(C)** Bar plot showing the cross-validation performance of IMMClock on the OneK1K dataset using intrinsic aging clocks for naïve/central memory (CM) and effector memory (EM) T cells, and abundance clocks. The Y-axis represents the Spearman correlation between immune age and chronological age at the person level. This panel showcases the performance of the individual modeling approaches described in (A), excluding the integrated model. **(D)** Bar plots depicting external validation for CD8⁺ T cell clocks on independent human datasets. The Y-axis represents the Spearman correlation between the predicted immune age and chronological age at the person level. All four types of clocks (intrinsic aging clocks for naïve/CM and EM T cells, abundance clocks, and the integrated clock) were evaluated across six independent human datasets: Terekhova et al. 2023^29^, Mogilenko et al. 2021^41^, Stephenson et al, 2021^40^, Perez et al. 2022^30^, AIDA^28^, and Lu et al. 2022^26^, left to right). **(E)** Bar plots depicting external validation for CD4⁺ T cell clocks on independent human datasets. Similarly to (D), all four types of clocks (intrinsic aging clocks for naïve/CM and EM T cells, abundance clocks, and the integrated clock) were evaluated across five independent human datasets: Terekhova et al. 2023^29^, Mogilenko et al. 2021^41^, Stephenson et al, 2021^40^, Perez et al. 2022^30^, AIDA^28^, left to right). The Y-axis represents the Spearman correlation between the predicted immune age and chronological age at the person level. Statistical notation: **** p < 0.0001; *** p < 0.001; ** p < 0.01; * p < 0.05; ns, not significant.

To study T cells immune aging more deeply, we next trained aging clocks specifically for naïve/central memory (CM) T cells and effector memory (EM) T cells to minimize the influence of compositional changes and to accurately capture intrinsic aging within these subtypes and to better capture the intrinsic aging processes in these populations. We next quantified age-related changes in cell composition by modeling the abundances of these T cell subtype to delineate how cell subtype proportions vary across different ages. Finally, we developed an integrated clock that includes these higher-resolution subtypes of T cells, to account for both the differentiation process and the intrinsic aging alterations. We stopped at this level and did not further refine cell type annotations, as previous studies^38,39^ have shown that higher-resolution distinctions are often difficult to make reliably, so we aimed to err on the side of caution.

We first quantified the effectiveness of IMMClock in predicting chronological age within the OneK1K cohort, using cross-validation for each immune cell type. Prediction accuracy was assessed by calculating the Spearman correlation between the predicted and actual ages across human samples. Specifically, we applied IMMClock to individual cells of a given immune cell type and aggregated the results by calculating the mean immune age of all relevant cells per person as the predicted age. As shown in **Supp. Fig. S1**, this method reassuringly produced similar results to applying the clock directly to pseudo-bulk data at the person level, affirming the performance and reliability of the clocks at individual cell resolution.

Among the four different IMMClock models generated for each of the five major immune subtypes, the integrated clocks demonstrated high accuracy for CD4⁺ T cells (Spearman ρ = 0.80, p-value=9.41e-218), CD8⁺ T cells (ρ = 0.68, p-value=6.6e-134), and NK cells (ρ = 0.56, p-value=1.4e-81) (**Fig. 2B**). The prediction accuracy was lower for B cells (ρ = 0.44, p-value=3.4e-48) and monocytes (ρ = 0.28, p-value=4.65e-19). These results align with previous findings that adaptive immune cells, such as CD4⁺ and CD8⁺ T cells, exhibit the most significant age-related changes^42^. Consequently, we focused all subsequent analyses on CD8⁺, CD4⁺ T cells and NK cells, for which IMMClock demonstrated robust prediction accuracy. As shown, the integrated IMMClock model, which accounts for both differentiation and intrinsic aging processes, achieved the highest overall accuracy. The naïve/CM-based IMMClock of both CD4⁺ and CD8⁺ T cells performed quite well, while the effector memory-based model demonstrated lower accuracy, as evaluated on the OneK1K dataset in cross-validation (**Fig. 2C**).

To evaluate the generalizability of IMMClock, we tested it on six independent PBMC single-cell datasets^26,28–30,40,41^, comprising 823 samples of CD8⁺ and CD4⁺ T cells from healthy donors (**Fig. 2D,E respectively**), without any additional model training. IMMClocks consistently demonstrated high predictive performance across cohorts (**Fig. 2D,E**, integrated clock in yellow), with Spearman correlations ranging from 0.56 to 0.88, underscoring its robustness across different cohorts and technologies. Further analysis revealed that the integrated clock maintained the highest predictive accuracy on external datasets, but the naïve/CM clock accuracy did not fall much behind, showing only a marginal difference (mean prediction accuracy: 0.72, 0.67 for integrated clock and naïve/CM clock respectively. p-value = 0.08, Wilcoxon test) (**Fig. 2D,E**). In contrast, the differentiation-based IMMClock, which relies solely on sub-cell-type proportions, is significantly less accurate than both the integrated (p-value = 0.0001, Wilcoxon test) and naïve/CM models (p-value = 0.004, Wilcoxon test), although it still outperformed the effector memory-based IMMClock (p-value = 0.02, Wilcoxon test). These findings show the fairly accurate performance of the integrated and the naïve/CM clocks in accurately estimating human’s biological age across many different independent datasets, for both CD8⁺ and CD4⁺ T cells.

We have demonstrated the effectiveness of IMMClock in predicting the immune age of individual cells, showing that the aggregated immune age across cells can robustly reflect the chorological age of individuals (see **Fig. 2B**). To further assess its predictive power at the individual cell level, we examined the association between immune age and key hallmarks of aging that can also be quantified from single-cell expression, specifically focusing on the activity of signatures relevant to telomere shortening, T cell senescence, and T cell exhaustion (**Fig. 3A–C)**. We focused on CD8⁺ T cells due to their well-characterized aging-related genes and signatures. Our results show that immune age is significantly correlated with the expression of genes involved in telomere shortening^42^ (e.g., *TERT*, *DKC1*), with T cell senescence (e.g., *KLRG1*^43^, *B3GAT1*^44^), and with T cell exhaustion^45^ (e.g., *LAG3*, *TIGIT*, *PDCD1*, *HAVCR2*, *CTLA4*) across individual cells, even after adjusting for chronological age. These results hold both in the OneK1K dataset and external datasets containing CD8⁺ T cells. Additionally, IMMClock reassuringly confirms that naïve CD8⁺ T cells have a younger immune age compared to memory CD8⁺ T cells (**Fig. 3D**), indicating that immune age increases along the T cell differentiation pathway. These analyses were conducted at the individual single-cell level, with each cell’s immune age adjusted by regressing out the donor’s chronological age and examining the residuals. The results are consistent with those observed at the donor level (**Supp. Fig. S2**).

**Figure 3.**
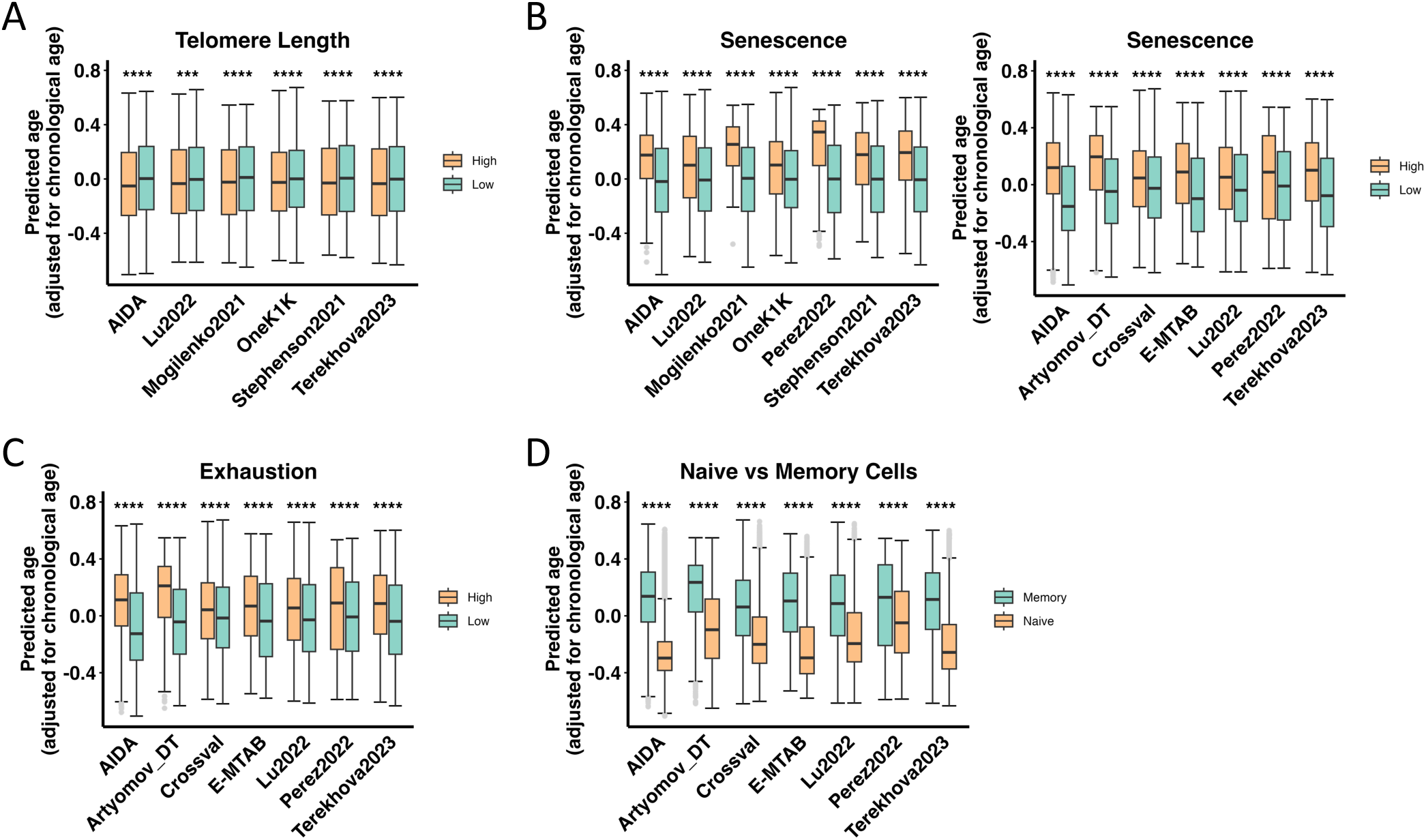
IMMClock associations with known CD8⁺ T cell signatures at single-cell resolution. This figure illustrates the association between predicted immune age and established T cell signatures of cellular phenotypes. We evaluated these associations using cross-validation on the OneK1K dataset and six independent scRNA-seq datasets: AIDA^28^, Mogilenko et al. 2021^41^, Terekhova et al. 2023^29^, Stephenson et al, 2021^40^, Perez et al. 2022^30^, and Lu et al. 2022^26^. The Y-axis represents the predicted immune age, adjusted for chronological age and sex. **(A)** Higher immune age is associated with reduced expression of telomere-maintenance genes (*TERT*, *DKC1*). **(B)** Increased immune age corresponds to higher expression of senescence-related genes (*KLRG1*, *B3GAT1*). **(C)** Higher immune age is linked to increased expression of exhaustion markers (*LAG3*, *TIGIT*, *PDCD1*, *HAVCR2*, *CTLA4*). **(D)** IMMClock estimates that naïve cells have a lower immune age compared to memory cells. Statistical notation (Mann-Whitney U test): **** p < 0.0001; *** p < 0.001; ** p < 0.01; * p < 0.05; ns, not significant.

### Cell type specificity and functional gene signatures enrichment of IMMClock

We next performed a comparative analysis of the three key immune cell-type IMMClock models —CD8⁺, CD4⁺, and NK cells—to evaluate their cell-type specificity. We evaluated the performance of each cell-type-specific model on expression data from the two other two cell types using a cross-validation approach. Reassuringly, the highest performance on each cell type is achieved by the model trained on that respective cell type compared to models trained on other cell types. (**Fig. 4A**). Furthermore, the genes selected as features by each cell-type-specific IMMClock are significantly enriched for gene markers associated with their respective cell types (Fisher’s test: p-values = 2e-16 for CD8⁺ T cells, 1.61e-7 for CD4⁺ T cells, and 2.46e-19 for NK cells; odds ratios = 15.42, 8.22, and 48.77, respectively) (see **Supp. Tables 2-8** for features, **Supp. Table 12** for cell-type-specific markers).

**Figure 4.**
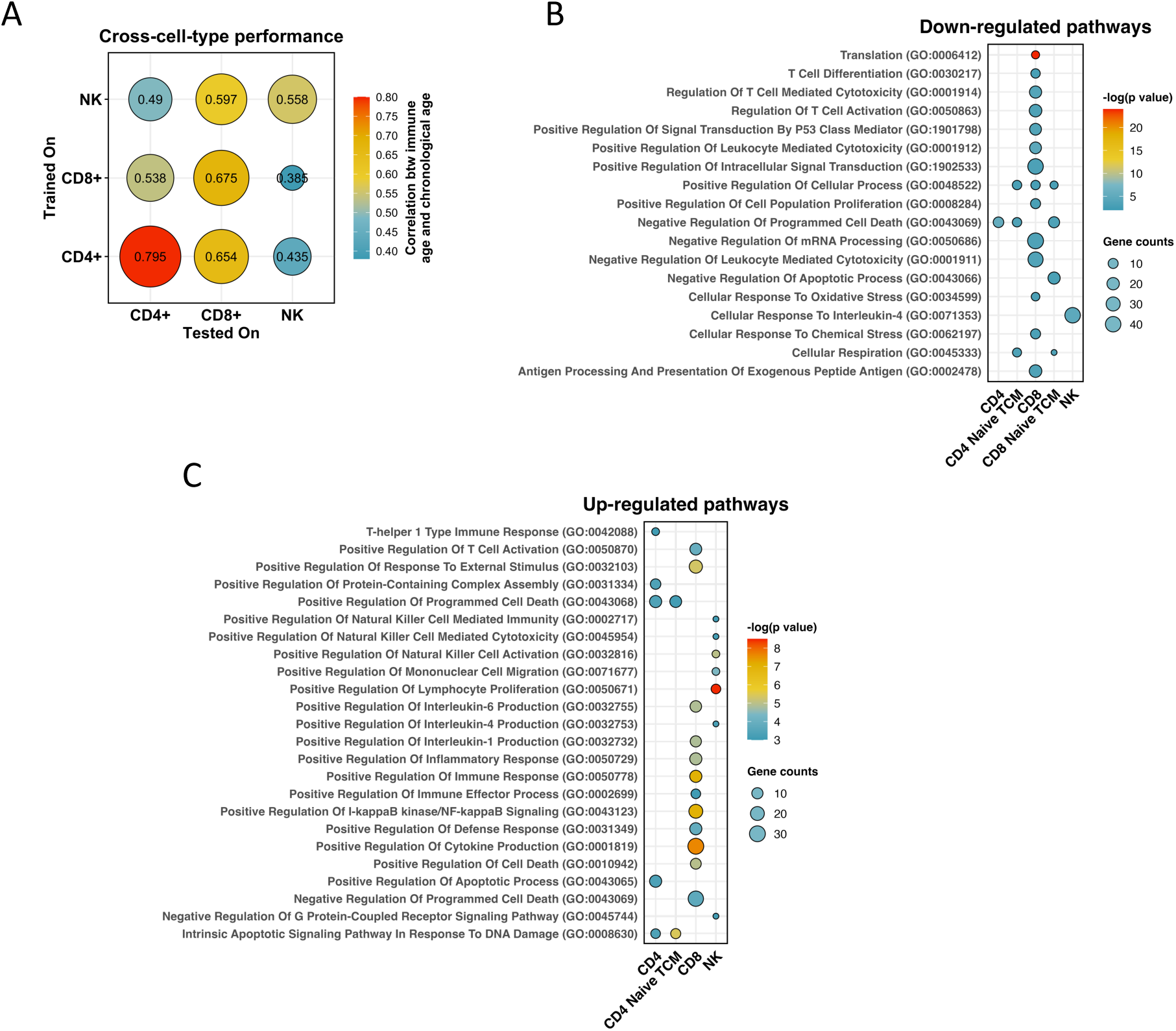
Specificity and functional analysis of cell-type-specific IMMClocks. **(A)** Evaluation of IMMClock’s cell type specificity through cross-model prediction using the OneK1K dataset. The relative size and color of the dots represent the Spearman correlation between predicted immune age and chronological age. **(B, C)** Enriched Gene Ontology (GO) biological processed among top feature genes selected by IMMClock, showing downregulated and upregulated pathways, respectively. The size of the dots corresponds to the number of overlapping genes, and the color represents the -log(p) value of the pathway.

We further explored the age-related characteristics captured by the IMMClock of each immune cell type, by conducting a pathway enrichment analysis of the genes selected as features in each cell-type-specific IMMClock model (**Fig. 4B, C**; see **Methods** for details on the analysis, **Supp. Tables 2-8** for the gene lists and **Supp. Tables 9, 10** for the enriched pathways). The downregulated pathways (**Fig. 4B**) provide notable insights into the mechanisms underlying immune aging. In CD8⁺ T cells, pathways related to the positive regulation of cellular processes, cell population proliferation, and translation are significantly downregulated, testifying to a decline in cellular growth, proliferation, and protein synthesis with aging. Additional downregulated pathways include negative regulation of leukocyte-mediated cytotoxicity, regulation of T cell activation, and T cell differentiation, indicating impaired cytotoxic functions, T cell responses, and differentiation processes, which are hallmarks of immune aging in CD8⁺ T cells^37,46^. In natural killer (NK) cells, there is a downregulation of cytokine response pathways, which are likely associated with decreased effective immune function of these cells as organisms age^47^. In CD4⁺ and CD8⁺ naïve and central memory T cells, downregulated pathways include regulation of apoptotic process, cellular respiration, and aerobic metabolism.

Conversely, many upregulated pathways (**Fig. 4C**) involve immune and inflammatory processes that become more active with aging. In CD8⁺ T cells, they include adaptive immune responses, T cell activation, cytokine-mediated signaling, and defense responses, which are often associated with chronic low-grade inflammation, also known as "inflammaging”^37^. Additional upregulated pathways involved in apoptosis and ER stress indicate that the IMMClock captures a heightened state of cellular damage and dysfunction (see **Supp. Table 9** for more details).

### Associations of IMMClock immune age estimations with different clinical phenotypes

We next turned to investigate whether cell-type-specific immune age is associated with specific clinical outcomes, after controlling for chronological age. We analyzed data from the Framingham Heart Study (FHS), a large-scale human cohort that provides comprehensive clinical and transcriptomic information from 2,691 healthy donors. Although it is a bulk gene expression cohort, we hypothesized that IMMClock can still be effectively applied, as we have demonstrated that the genes features utilized by IMMClock are predominantly cell type markers. Consequently, we first conducted tests that reassuringly revealed strong correlations between immune age estimations provided by the IMMClock (for CD4⁺ T cell, CD8⁺ T cell and NK cells) and the FHS study participants’ biological ages, achieving Spearman correlations of 0.57 for CD4⁺ T cells, 0.56 for CD8⁺ T cells, and 0.59 for NK cells (**Fig. 5A**). Additionally, immune age in CD4⁺ and CD8⁺ T cells is positively associated with C-reactive protein (CRP) levels, a well-established marker of inflammation^48^ (**Fig. 5B**). Notably, the IMMClocks, after adjusting for chronological age and sex, consistently show elevated immune age in FHS individuals with various diseases and adverse clinical conditions. A history of cancer, heart disease, and stroke is associated with a significantly heightened immune age across all three cell types (see **Fig. 5C, 5D, and 5F**). Using the Mann-Whitney U test, the p-values for cancer are 6.2e-03 for CD4⁺, 6e-04 for CD8⁺, and 3.9e-04 for NK cells. For stroke, the p-values are 1.4e-05 for CD4⁺, 1.4e-06 for CD8⁺, and 3.6e-04 for NK cells. In the case of heart disease, the p-values are 7.6e-05 for CD4⁺, 2.7e-07 for CD8⁺, and 2.7e-03 for NK cells.. Individuals with diabetes have significantly higher immune age in CD4⁺ and CD8⁺ T cells compared to controls (Mann-Whitney U test p-values: 8.3e-04 for CD4⁺ and 1.4e-02 for CD8⁺; see **Fig. 5E**). Intriguingly, deceased individuals (prior to 2019 as a cutoff point) had significantly higher immune age in CD4⁺ and CD8⁺ T cells compared to those still living (see **Fig. 5G**; Mann-Whitney U test p-values: 1.6e-02 for CD4⁺ and 2.2e-06 for CD8⁺). Further analysis in a systemic lupus erythematosus (SLE) dataset^30^ show that these patients have a significantly higher immune age in CD8⁺ T cells compared to healthy individuals (p-value: 6.6e-07) (**Fig. 5H**).

**Figure 5.**
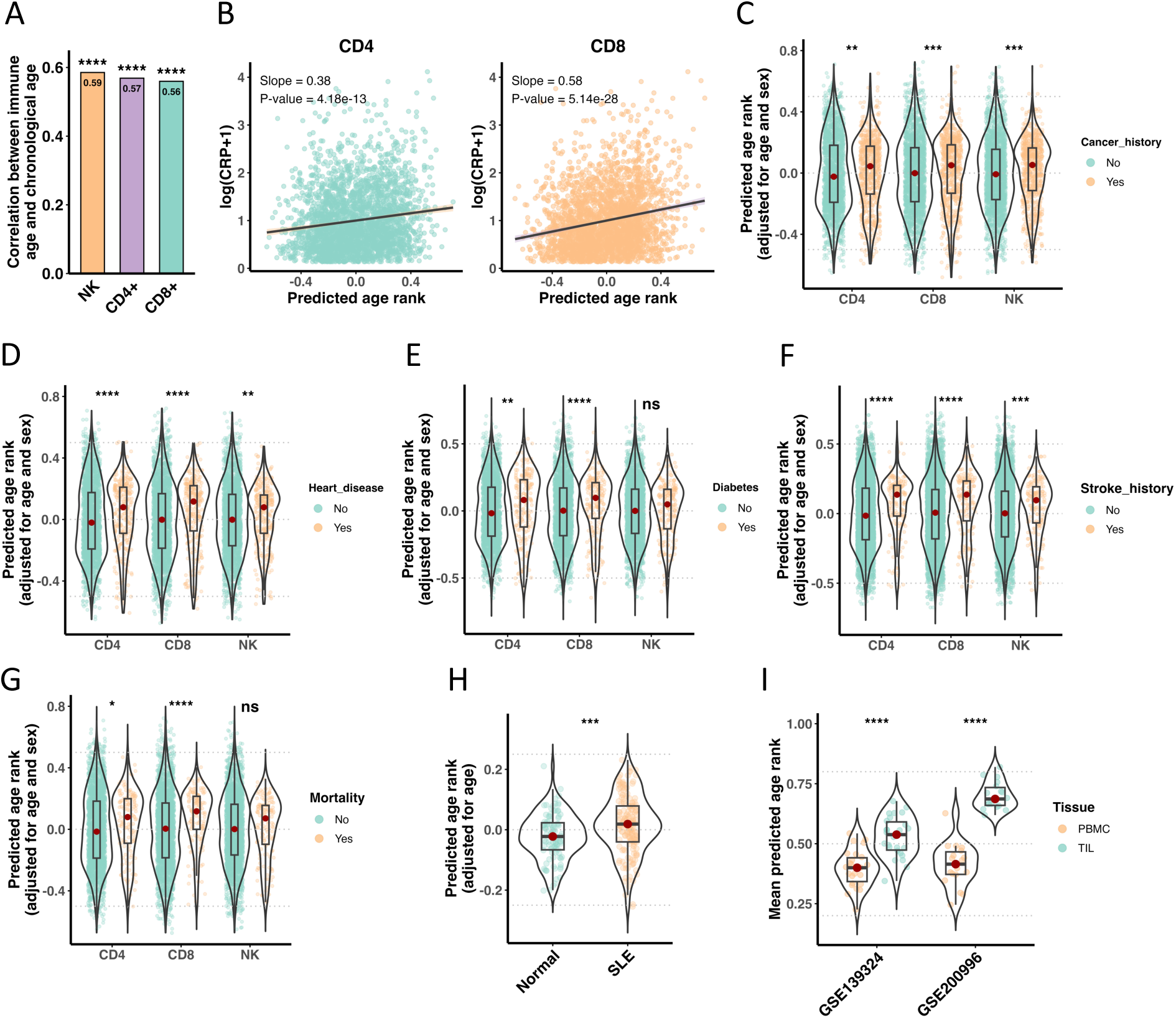
IMMClock associations with clinical phenotypes. This figure illustrates the relationships between immune age, as predicted by IMMClock, and various clinical phenotypes within the Framingham Heart Study dataset and other independent scRNA-seq cohorts. **(A)** A bar plot depicting IMMClocks predictive performance of chronological age in the Framingham Heart Study dataset. The Y-axis indicates correlation between predicted immune age and chronological age, using Spearman correlation, for each of the three main immune cell IMMClocks. **(B)** In the Framingham dataset, IMMClock age is significantly associated with CRP, an inflammatory marker; the X-axis represents predicted immune age, adjusted for study participant age. The Y-axis shows log(CRP+1). Trendline is the regression line of a linear regression, annotated are the slope and p-value. **(C-F)** In the Framingham dataset, individuals with a history of cancer (**C**), heart disease (**D**), diabetes (**E**), and stroke (**F**) exhibit higher immune age. **(G)** Individuals who died before 2019 had higher immune age at the time of blood sample collection compared to those who remained alive. **(H)** SLE (lupus) patients show higher immune compared to normal controls (data from Perez et al., 2022). The Y-axis shows the predicted immune age when adjusted by age. **(I)** In two HNSCC cohorts (Ruffin et al., 2021; Luoma et al., 2022), IMMClock indicates higher CD8⁺ immune age in tumor-infiltrating lymphocytes (TILs) within the tumor microenvironment (TME) compared to matched blood samples. Statistical notation: **** p < 0.0001; *** p < 0.001; ** p < 0.01; * p < 0.05; ns, not significant. Panels C-H: Mann-Whitney U test; Panel I: Wilcoxon test.

We next studied whether a previous history of cancer is associated with increased immune age across we applied IMMClock to two head and neck squamous cell carcinoma (HNSCC) cohorts^49,50^. These datasets are quite unique in that they contain single cell transcriptomics data from *matched* blood PBMC and tumor-infiltrating lymphocytes (TIL), enabling us to compare the immune age of circulating vs T cells in the tumor microenvironment (TME) *within the same patients.* Remarkably, we find that CD8⁺ TME TILs had a significantly higher immune age compared to circulating CD8⁺ T cells (p-value = 1.49e-08, p-value = 3.81e-06; **Fig. 5I**). CD4⁺ cells exhibited a similar trend (p-value = 0.08, p-value = 3.81e-06; see **Supp. Fig. S3**).

### IMMClock-inferred immune age is strongly associated with T cell functionality

We finally turned to study whether IMMClocks can assess the effects of gene perturbations on T cell functionality and predict functional outcomes based on immune age. To study this central question, we analyzed the dataset generated by Schmidt et al.^51^. In this investigation, Schmidt and colleagues aimed to identify regulators of T cell activation and cytokine production using CRISPR activation (CRISPRa) screens in primary human T cells. Primary human CD4^+^ and CD8^+^ T cells from two healthy donors were transduced with guide RNAs targeting 70 genes that included both positive and negative regulators of T cell activation (**Supp. Table 11**). The genetically perturbed T cells were either left *unstimulated (resting)* or *re-stimulated* via T cell receptor (TCR) engagement to mimic antigen-specific activation. Perturb-Seq Single-cell RNA sequencing was then performed to capture the gene expression profiles of individual T cells under each perturbation and condition. An *activation score* was then calculated by the original authors for the re-stimulated cells based on the expression levels of key activation-associated genes^51^. Using this score, the original authors quantified the functional response of T cells to stimulation, studying how each gene perturbation affected it. This dataset offered us a quite unique opportunity to systematically explore the relationship between immune aging and T cell functionality.

To this end, we applied IMMClocks to each of the 70 different CRISPRa Perturb-seq gene activation perturbations to investigate how they may have impacted immune age, and then study the latter’s relation to T cell functionality. Aiming to capture intrinsic aging related changes, we used the naïve/central memory (CM) clock to quantify immune age, given previous studies that have shown that T cell subsets early in differentiation exhibit the highest number of age-related gene expression changes, with naive and central memory cells being altered markedly more than effector memory cells^52–54^. Furthermore, naive and CM T cells are less influenced by prior immune activation and chronic inflammation compared to memory cells ^54^.

We first applied our clock to the NO-TARGET control cells—those without any gene perturbations. We observed that restimulated CD8⁺ T cells exhibit a significantly lower immune age compared to resting cells (**Fig. 6A**, p-value=3.35e-49). Next, we investigated the relationship between immune age in resting and restimulated cells across CRISPRa gene perturbations. We applied our clock at the individual cell level and aggregated the immune ages by calculating the mean cell immune age for each of the 70 gene perturbations. Plotting the mean immune ages of resting cells against those of restimulated cells for each gene perturbation shows a very strong correlation (Spearman *ρ* = 0.77, p-value=5.25e-15) between the immune ages of resting vs. restimulated cells across the different perturbations (**Fig. 6B**). This result suggests that while restimulation is associated with a shift toward a younger immune age at the population level, the relative age ranking of perturbations is largely preserved. Notably, certain perturbations, specifically those targeting negative regulators of stimulation-responsive cytokine production (e.g., *MUC1*, *MAP4K1*, *LAT2*), markedly deviate from this trend. In these cases, the restimulated cells exhibited higher immune ages than expected based on their resting state immune ages, suggesting that these gene perturbations influence immune age differently upon restimulation.

**Figure 6.**
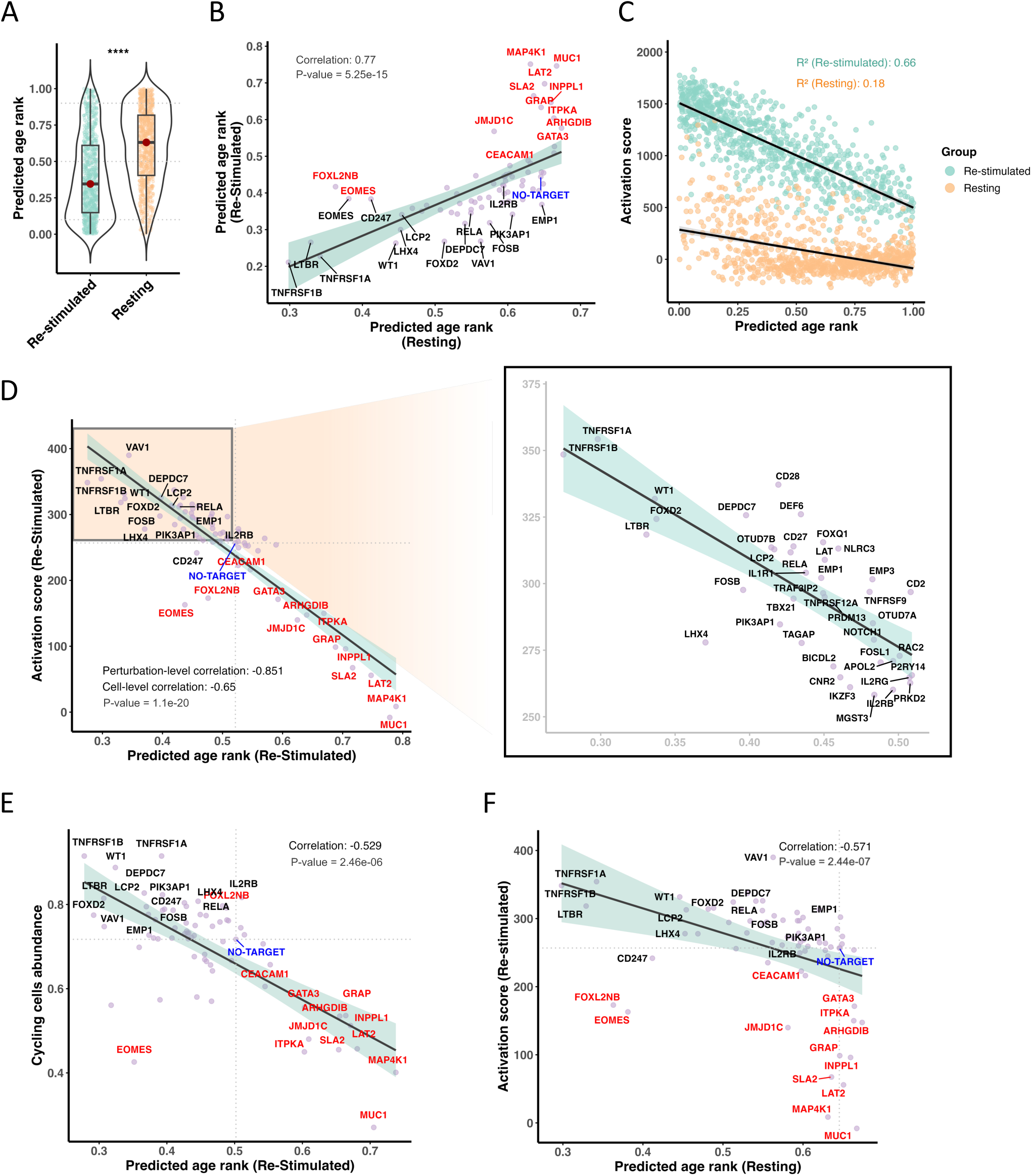
IMMClock reveals the effects of gene perturbations on CD8+ T cell immune age and activation. We applied the CD8 Naïve/TCM IMMClock to data from Schmidt et. al, 2022. **(A**) A violin plot demonstrating that restimulated unperturbed CD8⁺ T cells have significantly lower immune age compared to resting cells (based on NO-TARGET control cells without gene perturbations). The Y-axis is predicted immune age. **(B)** The immune age of stimulated cells is correlated with their age in the resting state across gene perturbations. The X-axis represents the predicted immune age of resting cells, and the Y-axis represents the predicted age of restimulated cells. Averaging immune age per perturbation across 70 gene perturbations revealed a strong correlation (ρ = 0.77), with certain perturbations (marked in red, e.g. *MUC1*, *MAP4K1*, *LAT2*) exhibiting higher immune age post-restimulation than expected given the overall trend. These are negative regulators of stimulation-responsive cytokine production^51^. Here and in the following panels, only a subset of the genes is labelled for visualization. **(C)** A scatter plot depicting the relationship between activation score (as inferred by the original authors^51^) and immune age as predicted by IMMClock, at the single-cell level. Based on NO-TARGET control cells without gene perturbations, predicted younger T cells exhibit higher activation scores; The X-axis shows predicted immune age of individual cells, and the Y-axis shows activation score. Restimulated cells are in green, and resting cells are in orange. **(D)** Across perturbations, there is a very strong negative correlation (*ρ*=-0.85) between the immune age of post-perturbation CD8+ T cells as predicted by IMMClock and their functional activation scores inferred in the original paper ^51^. Each marker point represents a gene perturbation. The gray dotted line indicates the immune age and activation score of unperturbed control cells (NO-TARGET), and the right panel zooms in on perturbations that result in younger and more activated cells. **(E)** X-axis is the predicted immune age of restimulated cells, but here we aggregate only G1 (non-cycling) phase cells for each perturbation. Y-axis is the relative proportion of cycling cells (S Phase, G2M Phase) within that perturbation. This panel shows that perturbations associated with a lower predicted immune age by IMMClock also increase the proportion of cycling cells**. (F)** X-axis is the predicted immune age of resting cells post-perturbation, and Y-axis is the activation score of stimulated cells post- perturbation (inferred by the original authors^51^). There is a strong negative correlation (ρ = −0.57) between the predicted immune age of resting cells post-perturbation and the activation scores of stimulated cells post-perturbation across all perturbations. This indicates that IMMClock can identify perturbations leading to younger and more activated cells even before stimulation.

To investigate the fundamental relationship between immune age and T cell activation, we first plotted the activation scores of individual cells against immune age in the unperturbed, NO-TARGET case. In the case of restimulated cells, this revealed a strong inverse relationship between the two, as indicated by a negative slope and an R- squared (R^2^) of 0.66 (**Fig. 6C**). To study this relation across perturbations, we aggregated the mean immune age and activation scores of the individual cells for each of the 70 gene perturbations. Analogous to our previous observations, we identify a very strong negative correlation (*ρ*= −0.85, p-value=1.1e-20) between immune age and activation score (**Fig. 6D**). This finding testifies that perturbations leading to younger T cells are also strongly associated with higher activation levels. The top perturbations that generate cells that are both younger and highly activated denote potential targets for enhancing T cell functionality. Importantly, this negative correlation remains robust even when controlling for cell cycle phases by analyzing only G1 phase cells **(see Supp. Fig. S4**). Notably, the genes in the CD8 naïve/TCM clock show no significant overlap with those used by the original authors to infer the activation score (odds ratio=0.69, p-value=0.99, Fisher’s exact test, see **Methods**), ruling out the possibility that the strong correlation between immune age and activation scores arises trivially from their potential overlap. Interestingly, two perturbations, *EOMES* and *FOXL2NB*, result in cells with younger immune ages (p- value=3.7e-10, p-value=3.16e-15, respectively) but lower activation levels (p- value=2.75e-6, p-value=4.43e-11, respectively) than the control. These gene perturbations may decouple immune age from activation, potentially influencing other aspects of T cell biology such as differentiation or suppressive functions.

Recognizing the close relationship between activation and proliferation, we further investigated the potential role of proliferation as a key factor associated with T cell activation. In the NO-TARGET control cells, CD8⁺ T cells in the cycling phases (S/G₂M) exhibited a younger immune age compared to those in the G₁ phase (**see Supp. Fig. S5**). Notably, gene perturbations that result in younger immune ages of cells at the G₁ phase also increase the proportion of cycling cells (**Fig. 6E**). These findings suggest that there possibly may be an interplay between T cell proliferation and immune aging.

Finally, we asked whether the immune age of *resting* post-perturbation cells could predict the magnitude of their activation response upon antigen exposure. By plotting the immune age of resting cells against their activation scores after restimulation across all gene perturbations, we find that the immune age of resting cells is a strong predictor of their subsequent re-stimulation activation levels (**Fig. 6F**). This suggests that immune age, as determined in the resting state, can reliably predict the functional responsiveness of T cells following stimulation, without the need for additional stimulation and functional assays. Overall, our findings demonstrate that gene perturbations that promote enhanced activation are generally associated with the emergence of younger T cells with increased proliferation. This highlights the potential of targeting specific genes that rejuvenate T cells to bolster immune responses.

## Discussion

In this study we introduce IMMClock, a single-cell resolution transcriptomic aging clock designed to quantify immune age at both person and single-cell levels. IMMClock was successfully validated across diverse independent single-cell datasets. It can effectively quantify the effects of gene perturbations on T cell functionality, identifying interventions that reduce immune age and result in activated T cells. Understanding how specific genetic perturbations can modulate immune aging and restore T cell efficacy is a pivotal frontier in immunology with considerable therapeutic potential.

IMMClock enables person-level and cell-level estimation of immune age. As a person-level clock, IMMClock exhibits strong predictive accuracy for immune age in CD4, CD8 T cells and NK cells, which was validated across multiple independent cohorts, providing a view of aging of individual immune cell populations. At the single-cell level, IMMClock highlights associations between immune age and established cellular processes, such as senescence, exhaustion, and telomere shortening. A central finding of our study is that gene perturbations that promote T cell activation and proliferation also mostly lead to younger T cells, underscoring the link between T cell functionality and immune aging.

IMMClock builds upon and advances previous efforts in developing cell-type-specific aging clocks. Earlier studies have primarily focused on aging clocks based on DNA methylation patterns^21,22^, cell-type abundances^11–13^, or have been conducted using mouse models^24^. Notably, Lu et al.^26^ and Buckley et al.^24^ pioneered the first cell-type-specific aging clocks for CD8⁺ T cells and neurons, respectively. Going beyond these foundational studies, IMMClock make a substantial contribution to this field by being trained and tested on large-scale human datasets, effectively controlling for changes in cell type composition, and being applicable to both single-cell and bulk transcriptomics data. At the individual single-cell level, the previous development of aging clocks has been quite limited, with only a few studies such as scEpiAge (focused on methylation^22^) and Lu et al.^26^ making notable strides. IMMClock makes further advances by providing a high-resolution, single-cell immune age assessment tool, offering deeper insights into the aging processes of individual immune cells and enhancing their potential use for precise immunological interventions.

Our study obviously has several limitations. First, IMMClock is trained on cross-sectional data, limiting its capacity to track immune age changes within individuals over time. Longitudinal data would provide a more accurate understanding of immune aging dynamics throughout the lifespan. Unfortunately, high-resolution longitudinal single-cell datasets are currently scarce. Second, within subsets like the naïve CD8+ T cells population, there is substantial diversity. The inherent heterogeneity within T cell populations poses challenges for single-cell annotations^55^. Evidently, aging-related transcriptional changes can obscure cell identity^56^, making it difficult to refine annotations accurately, which remains an important research challenge on its own.

Future research should aim to validate IMMClock findings in *in vivo* models to assess the therapeutic potential of identifying gene perturbations that concomitantly make T cells both younger and more activated in more physiologically relevant settings. Expanding the range of genes that are screened and tested in such activation studies will provide a more comprehensive understanding of the genetic regulation of T cell aging and identify additional candidates for immunomodulation. Integrating IMMClock with other omics data, such as proteomics and epigenomics, could further refine the immune age clock and uncover deeper mechanistic insights into T cell aging. Additionally, exploring the use of IMMClock in predicting the levels of specific immune-related functions, such as vaccine responses and cancer immunotherapy outcomes, could broaden its applicability and impact in clinical settings.

In conclusion, IMMClock represents a powerful and versatile tool offering new insights into the interplay between immune cell functionality and aging. Our findings indicate that targeting specific genes that enhance T cell activation and proliferation usually also leads to a younger T cell population, which may possibly further contribute to improving immune responses on extended time scales. By elucidating the intricate relationships between immune aging, cell activation, and cell cycle dynamics, this study lays the groundwork for developing targeted immunotherapies aimed at concomitantly mitigating immune senescence and enhancing overall immune competence. Future applications of IMMClock in clinical settings hold promise for identifying and validating interventions that both "rejuvenate" and activate T cells based on their transcriptomic profiles, enhancing experimental efforts to slow down and counteract age-related immune decline.

## Methods

### Preprocessing of the OneK1K dataset and other single-cell datasets

We obtained the OneK1K PBMC single cell data from Yazar et al., 2022^27^ as our training datasets, along with six additional PBMC single-cell datasets^26,28–30,40,41^ for validation. For the OneK1K cohort, we utilized the cell type annotations provided in the original study, while we reannotated the six additional validation single-cell datasets to ensure the consistent cell type classifications. This was accomplished using *CellTypist*^38^ with the Immune_All_Low.pkl model.

Given the diversity of cell types, we focus on five major immune cell types—CD4⁺ T cells, CD8⁺ T cells, monocytes, NK cells, and B cells. For datasets with read counts, we derived Counts Per Million (CPM) values by normalizing the read counts based on library size, followed by a logarithmic transformation to stabilize variance for downstream analyses. For datasets that had already undergone the same normalization in the original studies, we utilized those values directly.

To further prepare the OneK1K dataset for training our aging clocks, we aggregated individual cells into cell type-specific pseudo-bulk samples for each person. We implemented uniformed quality control across all the single cell datasets, excluding cells with fewer than 200 expressed genes, genes detected in fewer than three cells, and cells exhibiting over 5% mitochondrial gene expression to maintain data integrity. All preprocessing steps were performed using the *Scanpy*^57^ library in Python.

### The IMMClock Model

1. **Preprocessing**: we standardized the gene expression values, which serve as feature values, by applying z-score transformation. We performed a Box-Cox transformation on the chronological ages, serving as the target labels. This transformation aligns the age distribution more closely with normality and corrects any biases present in the original age distribution, enhancing the model’s predictive capabilities.
2. **Feature (gene) selection:** to reduce feature space and enhance the prediction power of our model, we performed feature selection before training. We identified genes significantly (p < 0.01 after Bonferroni correction) associated with chronological age. We further refine our gene selection by choosing the top 50% of genes most positively correlated with age and the top 50% most negatively correlated with age. Importantly, this feature selection step is performed exclusively on the training dataset to prevent data leakage.
3. **Dimensionality reduction using PCA:** We applied principal component analysis (PCA) to the selected genes, preserving 80% of the explained variance. The PCA transformation was performed exclusively on the training dataset and is later applied to the test or validation dataset.
4. **Model training**: We aim to predict the chorological age by leveraging the generated principal components as features for each cell type across persons. The explicit model is represented by the following equation:

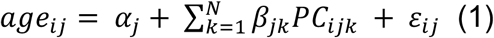

In this equation, *age_ij_*_"_ represents the chorological age for *i*th individual and *j*th cell type (e.g., CD4^+^ T cells, CD8^+^ T cells and B cells). Here, *±_j_* serves as the baseline age level for *j*th cell type, *PC_ijk_* denotes the *K*th principal components (PC) out of the total *N* PCs for *i*th individual and *j*th cell type, *²_jk_* is the coefficient for the *K*th PC and *j*th cell type, and *ε_ij_* represents the residuals for *i*th individual and *j*th cell type.

To further reduce overfitting and enhance generalization, we employed elastic net linear model^58^ from the *scikit-learn*^59^ library for training. First, stratified 5-fold cross-validation was performed to identify the optimal hyperparameters, lambda and alpha, which achieves the optimal performance in a cross-validation manner. Following that, the final training was conducted using the optimized hyperparameters to enable independent validation and immune age prediction given unseen samples or individual cells.

**5. Performance assessment metrics:** To assess the model’s performance, we calculate the Spearman correlation between the percentile ranks of actual ages and the percentile ranks of the predicted ages. This non-parametric measure offers a robust assessment of the model’s performance.

### Technical validation of IMMClock

We performed both cross-validation and independent validation to assess IMMClock’s performance in predicting chronological ages. For cross-validation, we employed a repeated stratified 5-fold approach using scikit-learn^59^ in Python, specifically utilizing the function RepeatedStratifiedKFold(). To minimize sampling bias, we performed 3 repeats with stratification to ensure each fold maintains a similar age distribution. For independent validation, we assessed the model’s robustness and generalizability to new, unseen datasets, by training it on the entire OneK1K dataset and validate it on all other independent single cell datasets. When training for each independent validation, we used only the intersection of genes present in both the OneK1K dataset and the independent validation dataset.

We applied IMMClock independently to each immune cell type within the OneK1K dataset. For each cell type, the model was trained on cell type-specific pseudo-bulk samples derived from the training group. The trained model then infers an immune age for each individual immune cell. To derive an immune age for an individual, we compute the mean immune age of all the immune cells from that specific cell type. We also calculate performance metrics, including Spearman correlation coefficients and mean squared error (MSE) between the percentile ranks of actual and predicted ages, to assess predictive accuracy.

### Biological validation of IMMClock

To further validate IMMClock, we measure the association between our inferred immune ages and relevant gene signatures, including markers for telomere length, T cell senescence, immune cell differentiation, and exhaustion. All these tests are performed at both the individual cell and person levels for each cell type. At the individual cell level, we adjusted the predicted immune age of each cell by the chronological age of the individual using a linear regression model. Specifically, the linear regression model treats the predicted immune age as the dependent variable and the chronological age as the independent variable. The residuals from this regression, representing the portion of immune age not explained by chronological age, were used as the adjusted immune age values. This adjustment controls for the confounding effect of chronological age, ensuring that the associations between gene expression and immune age are not influenced by individuals’ chronological age. Subsequently, cells are categorized to high-expression-level (> 0) and low-expression-level groups (=0) based on specific gene expression signature. We then compared the adjusted immune ages between these groups using the Mann-Whitney U test. At the person level, for each specific gene signature, we first divide cells to two groups, the high-expression cells and the low-expression cells for each individual based on the expression level of that gene signature. Next, we calculate the average immune age for each group, resulting in two immune ages per person – one for the high-expression group and one for the low-expression group. We then performed a paired Wilcoxon signed-rank test to compare these two groups across individuals. Since comparisons are made within the same person, this approach inherently accounts for individual age differences, eliminating the need for additional age adjustments.

### Cell-Type Specificity Analysis

To evaluate the cell-type specificity of the IMMClock models for CD4⁺ T cells, CD8⁺ T cells, and NK cells, we conducted a gene enrichment analysis comparing the genes selected as features by each cell-type-specific IMMClock to established cell-type-specific markers from CellMarker 2.0^60^. Using Fisher’s exact test, we determined that the genes selected by each IMMClock were significantly enriched for markers associated with their respective cell types.

### Functional Analysis of IMMClock-Related Genes

To assess the contribution of each gene in the prediction of chronological age, we extracted gene weights/coefficients derived from the elastic net model. To account for the dimensionality reduction performed by PCA analysis, we integrated these coefficients with the PCA components. Specifically, we calculated the overall weight for each gene by projecting the elastic net coefficients back into the original gene space through a matrix multiplication with the PCA components.

To perform a functional analysis of genes relevant to IMMClock, we utilized the Enrichr webserver interface^61^. We selected genes identified by the main IMMClocks across the following cell types: CD8⁺, CD8 Naive, CD8 Naive TCM, CD4⁺, CD4 Naive, CD4 Naive TCM, CD4 TEM, CD8 TEM, and NK cells. These genes were categorized based on their weights in the clock models: genes with positive weights, indicating a positive effect on immune age, and genes with negative weights, indicating a negative effect on immune age. Each of these gene groups was separately input into Enrichr to identify enriched Gene Ontology Biological Process (GO_BP) terms using the "GO_Biological_Process_2023" library. We applied a false discovery rate (FDR) threshold of <0.05 to determine statistical significance. To focus on pathways relevant to immune cell functions, we further filtered the enriched pathways by retaining those containing specific keywords, including *regulation, inflammation, immune, T cell, signaling, antigen, stimulus, activation, differentiation, proliferation, chemotaxis, cellular response, migration, adhesion, interferon, cytokine, cytotoxicity, apoptosis, interleukin, production, and translation*.

### Framingham Heart Study Data Analysis

Access to various parts of the Framingham Heart Study (FHS) data was obtained via human subject protocol 10-HG-N045 approved by an Institutional Review Board (IRB) at the National Institutes of Health and via permission from the database of Genotypes and Phenotypes (dbGaP). Normalized RNA-seq gene expression data and phenotypes were downloaded from dbGaP (study identifier phs000794, phs000007) and include the individuals from the Offspring and Generation 3 cohorts who attended the ninth and second clinical exams, respectively, and who signed either HMB-IRB- MDS or HMB-IRB- NPU-MDS research consent forms (*n* = 720 and 1,971 for Offspring and Generation 3, respectively).

To evaluate the association between immune age and various phenotypic traits within the FHS cohort, we first adjusted the predicted immune ages to account for the confounding effects of chronological age and sex, as discussed in the “biological validation of IMMClock” subsection of the **Methods**. These adjusted immune ages were then utilized for the comparative analyses across different phenotypic states, including diabetes, cancer, heart disease, stroke, mortality and smoking history. For instance, we compared the distribution of adjusted immune ages between individuals diagnosed with diabetes and those without, allowing for the assessment of associations between immune aging and disease status while controlling for demographic variables.

### Analysis of gene perturbations and their effects on CD8^+^ T cell immune age and activation

We obtained the perturb-seq dataset from the Schmidt et al.’s study^51^ via Zenodo (https://zenodo.org/records/5784651). We loaded the dataset into Python using the *h5py*^62^ library and utilized the normalized data from the original study using *SCTransform*^63^ for our downstream analysis. Additionally, we leveraged the original cell-type annotations for CD4⁺/CD8⁺ T cells, inferred activation scores, and cell cycle phase annotations. For this analysis, we employed the CD8+ Naïve/TCM IMMClock. First, we isolated the unperturbed control cells (NO-TARGET) and applied IMMClock directly to both resting and restimulated cells to estimate their immune ages and perform the comparison (see Fig. 6 and 7). Second, we evaluated the effect of the 70 different gene perturbations on immune age by applying IMMClock to the relevant cells. The averaged immune ages were computed across cells for each gene perturbation to derive the assessment at the perturbation level. Following that, we assess the association between the predicted immune age and activation levels across perturbations using Spearman correlation analysis (see Fig. 6B and 6D-F). Likewise, we applied the same analysis to individual single cells to evaluate the association between the immune age and activation cells across cells.

## Data availability

All datasets used in this study are from publicly available sources, but the Framingham Heart Study data are different from the other data sets in that access requires permission from an Institutional Review Board and from dbGaP. Links to download the datasets can be found in **Supp. Table 1**.

## Code availability

The code for reproducing the results of this study will be made available for academic research purposes after publication via Zenodo.

## Author Contributions

Conceptualization of the study: Y.G.S., K.W. and E.R. Data collection and curation: Y.G.S. with assistance from K.W., A.A.S., S.M., S. Sinha and S. Sahni. Selection of methods for data analysis: Y.G.S. and K.W., with assistance from V.G., S.R.D., B.W. Visualization: Y.G.S. and D.W., with assistance from K.W. Literature search of related work: Y.G.S. and K.W., with assistance from S.M. Writing first draft: Y.G.S, K.W. and E.R. Writing later drafts and editing: Y.G.S., K.W., E.R., A.A.S., N.P.R. and N.W.

Supervised the study: K.W. and E.R.

## Acknowledgements

This research is supported in part by the Intramural Research Program of the National Institutes of Health, National Cancer Institute, Center for Cancer Research. This work utilized the computational resources of the NIH HPC Biowulf cluster (http://hpc.nih.gov). Fig. 1 panel C, Fig. 2 panel A created with BioRender.com. We acknowledge Rémy Bosselut, Avinash Bhandoola, and Sridhar Hannenhalli for valuable discussions and insightful input, which greatly contributed to the refinement of this work.

## References

1. Rutledge, J., Oh, H. & Wyss-Coray, T. Measuring biological age using omics data. Nat Rev Genet 23, 715–727 (2022).

2. Hannum, G. et al. Genome-wide Methylation Profiles Reveal Quantitative Views of Human Aging Rates. Mol Cell 49, 359–367 (2013).

3. Horvath, S. DNA methylation age of human tissues and cell types. Genome Biol 14, R115 (2013).

4. Lu, A. T. et al. DNA methylation GrimAge strongly predicts lifespan and healthspan. Aging 11, 303–327 (2019).

5. Levine, M. E. et al. An epigenetic biomarker of aging for lifespan and healthspan. Aging 10, 573– 591 (2018).

6. Peters, M. J. et al. The transcriptional landscape of age in human peripheral blood. Nat Commun 6, 8570 (2015).

7. Fleischer, J. G. et al. Predicting age from the transcriptome of human dermal fibroblasts. Genome Biol 19, 221 (2018).

8. Wang, K. et al. Comprehensive map of age-associated splicing changes across human tissues and their contributions to age-associated diseases. Sci Rep 8, 10929 (2018).

9. Chatsirisupachai, K., Palmer, D., Ferreira, S. & de Magalhães, J. P. A human tissue-specific transcriptomic analysis reveals a complex relationship between aging, cancer, and cellular senescence. Aging Cell 18, (2019).

10. Meyer, D. H. & Schumacher, B. BiT age: A transcriptome-based aging clock near the theoretical limit of accuracy. Aging Cell 20, (2021).

11. Sayed, N. et al. An inflammatory aging clock (iAge) based on deep learning tracks multimorbidity, immunosenescence, frailty and cardiovascular aging. Nat Aging 1, 598–615 (2021).

12. Alpert, A. et al. A clinically meaningful metric of immune age derived from high-dimensional longitudinal monitoring. Nat Med 25, 487–495 (2019).

13. Sparks, R. et al. A unified metric of human immune health. Nature Medicine 2024 30:*9* 30, 2461–2472 (2024).

14. Hung, K. et al. The Central Role of CD4+ T Cells in the Antitumor Immune Response. Journal of Experimental Medicine 188, 2357–2368 (1998).

15. Raskov, H., Orhan, A., Christensen, J. P. & Gögenur, I. Cytotoxic CD8+ T cells in cancer and cancer immunotherapy. BriDsh Journal of Cancer 2020 124:*2* 124, 359–367 (2020).

16. Gruver, A. L., Hudson, L. L. & Sempowski, G. D. Immunosenescence of ageing. J Pathol 211, 144– 156 (2007).

17. Weiskopf, D., Weinberger, B. & Grubeck-Loebenstein, B. The aging of the immune system. Transplant InternaDonal 22, 1041–1050 (2009).

18. Pawelec, G. Age and immunity: What is “immunosenescence”? Exp Gerontol 105, 4–9 (2018).

19. Bektas, A., Schurman, S. H., Sen, R. & Ferrucci, L. Human T cell immunosenescence and inflammation in aging. J Leukoc Biol 102, 977–988 (2017).

20. Crooke, S. N., Ovsyannikova, I. G., Poland, G. A. & Kennedy, R. B. Immunosenescence and human vaccine immune responses. Immunity & Ageing 2019 16:*1* 16, 1–16 (2019).

21. Trapp, A., Kerepesi, C. & Gladyshev, V. N. Profiling epigenetic age in single cells. Nature Aging 2021 1:*12* 1, 1189–1201 (2021).

22. Bonder, M. J. et al. scEpiAge: an age predictor highlighting single-cell ageing heterogeneity in mouse blood. Nature CommunicaDons 2024 15:*1* 15, 1–15 (2024).

23. Weng, N. Transcriptome-based measurement of CD8+ T cell age and its applications. Trends Immunol 44, 542–550 (2023).

24. Buckley, M. T. et al. Cell-type-specific aging clocks to quantify aging and rejuvenation in neurogenic regions of the brain. Nat Aging 3, 121–137 (2022).

25. Zhu, H. et al. Human PBMC scRNA-seq–based aging clocks reveal ribosome to inflammation balance as a single-cell aging hallmark and super longevity. Sci Adv 9, (2023).

26. Lu, J. et al. Heterogeneity and transcriptome changes of human CD8+ T cells across nine decades of life. Nat Commun 13, 5128 (2022).

27. Yazar, S. et al. Single-cell eQTL mapping identifies cell type-specific genetic control of autoimmune disease. Science (1979) 376, (2022).

28. Kock, K. H. et al. Single-cell analysis of human diversity in circulating immune cells. *bioRxiv* 20, 2024.06.30.601119 (2024).

29. Terekhova, M. et al. Single-cell atlas of healthy human blood unveils age-related loss of NKG2C+GZMB−CD8+ memory T cells and accumulation of type 2 memory T cells. Immunity 56, 2836–2854.e9 (2023).

30. Perez, R. K. et al. Single-cell RNA-seq reveals cell type-specific molecular and genetic associations to lupus. Science (1979) 376, (2022).

31. Liu, C. et al. Whole genome DNA and RNA sequencing of whole blood elucidates the genetic architecture of gene expression underlying a wide range of diseases. Sci Rep 12, 20167 (2022).

32. Taliun, D. et al. Sequencing of 53,831 diverse genomes from the NHLBI TOPMed Program. Nature 590, 290–299 (2021).

33. Tong, H. et al. Cell-type specific epigenetic clocks to quantify biological age at cell-type resolution. bioRxiv 2024.07.30.605833 (2024) doi:10.1101/2024.07.30.605833.

34. Tomusiak, A. et al. Development of an epigenetic clock resistant to changes in immune cell composition. CommunicaDons Biology 2024 7:*1* 7, 1–13 (2024).

35. Goronzy, J. J., Fang, F., Cavanagh, M. M., Qi, Q. & Weyand, C. M. Naive T cell maintenance and function in human aging. J Immunol 194, 4073–4080 (2015).

36. Mogilenko, D. A., Shchukina, I. & Artyomov, M. N. Immune ageing at single-cell resolution. Nat Rev Immunol 22, 484–498 (2022).

37. Miqelbrunn, M. & Kroemer, G. Hallmarks of T cell aging. Nature Immunology 2021 22:*6* 22, 687–698 (2021).

38. Domínguez Conde, C., et al. Cross-tissue immune cell analysis reveals tissue-specific features in humans. Science (1979) 376, (2022).

39. Andreaqa, M. et al. Interpretation of T cell states from single-cell transcriptomics data using reference atlases. Nature CommunicaDons 2021 12:*1* 12, 1–19 (2021).

40. Stephenson, E. et al. Single-cell multi-omics analysis of the immune response in COVID-19. Nat Med 27, 904–916 (2021).

41. Mogilenko, D. A. et al. Comprehensive Profiling of an Aging Immune System Reveals Clonal GZMK+ CD8+ T Cells as Conserved Hallmark of Inflammaging. Immunity 54, 99–115.e12 (2021).

42. Armanios, M. & Blackburn, E. H. The telomere syndromes. Nat Rev Genet 13, 693 (2012).

43. Henson, S. M. & Akbar, A. N. KLRG1-more than a marker for T cell senescence. Age (Omaha) 31, 285–291 (2009).

44. Brenchley, J. M. et al. Expression of CD57 defines replicative senescence and antigen-induced apoptotic death of CD8+ T cells. Blood 101, 2711–2720 (2003).

45. van der Leun, A. M., Thommen, D. S. & Schumacher, T. N. CD8+ T cell states in human cancer: insights from single-cell analysis. Nature Reviews Cancer 2020 20:*4* 20, 218–232 (2020).

46. Zhang, H., Weyand, C. M. & Goronzy, J. J. Hallmarks of the aging T-cell system. FEBS J 288, 7123– 7142 (2021).

47. Brauning, A. et al. Aging of the Immune System: Focus on Natural Killer Cells Phenotype and Functions. Cells 2022*, Vol.* 11, *Page* 1017 **11**, 1017 (2022).

48. Sproston, N. R. & Ashworth, J. J. Role of C-reactive protein at sites of inflammation and infection. Front Immunol 9, 342848 (2018).

49. Ruffin, A. T., et al. B cell signatures and tertiary lymphoid structures contribute to outcome in head and neck squamous cell carcinoma. Nat Commun 12, 3349 (2021).

50. Luoma, A. M. et al. Tissue-resident memory and circulating T cells are early responders to pre-surgical cancer immunotherapy. Cell 185, 2918–2935.e29 (2022).

51. Schmidt, R. et al. CRISPR activation and interference screens decode stimulation responses in primary human T cells. Science (1979) 375, (2022).

52. Thomson, Z. et al. Trimodal single-cell profiling reveals a novel pediatric CD8αα+ T cell subset and broad age-related molecular reprogramming across the T cell compartment. Nature Immunology 2023 24:*11* 24, 1947–1959 (2023).

53. Moskowitz, D. M. et al. Epigenomics of human CD8 T cell differentiation and aging. Sci Immunol 2, 17 (2017).

54. Gong, Q. et al. Longitudinal Multi-omic Immune Profiling Reveals Age-Related Immune Cell Dynamics in Healthy Adults. bioRxiv 2024.09.10.612119 (2024) doi:10.1101/2024.09.10.612119.

55. Mullan, K. A., de Vrij, N., Valkiers, S. & Meysman, P. Current annotation strategies for T cell phenotyping of single-cell RNA-seq data. Front Immunol 14, 1306169 (2023).

56. Connolly, E. et al. Loss of immune cell identity with age inferred from large atlases of single cell transcriptomes. Aging Cell 00, e14306 (2024).

57. Wolf, F. A., Angerer, P. & Theis, F. J. SCANPY: large-scale single-cell gene expression data analysis. Genome Biol 19, 15 (2018).

58. Zou, H. & Hastie, T. Regularization and Variable Selection Via the Elastic Net. J R Stat Soc Series B Stat Methodol 67, 301–320 (2005).

59. Pedregosa, F. et al. Scikit-learn: Machine Learning in Python. the Journal of machine Learning research 12, 2825–2830 (2011).

60. Hu, C. et al. CellMarker 2.0: an updated database of manually curated cell markers in human/mouse and web tools based on scRNA-seq data. Nucleic Acids Res 51, D870–D876 (2023).

61. Xie, Z. et al. Gene Set Knowledge Discovery with Enrichr. Curr Protoc 1, (2021).

62. Colleqe, A. Python and HDF5: Unlocking ScienDfic Data. (O’Reilly Media, Inc, 2013).

63. Choudhary, S. & Satija, R. Comparison and evaluation of statistical error models for scRNA-seq. Genome Biol 23, 1–20 (2022).

