## Supplemental Figures for "IMMClock reveals immune aging and T cell function at single-cell resolution"

### Supplementary Figures

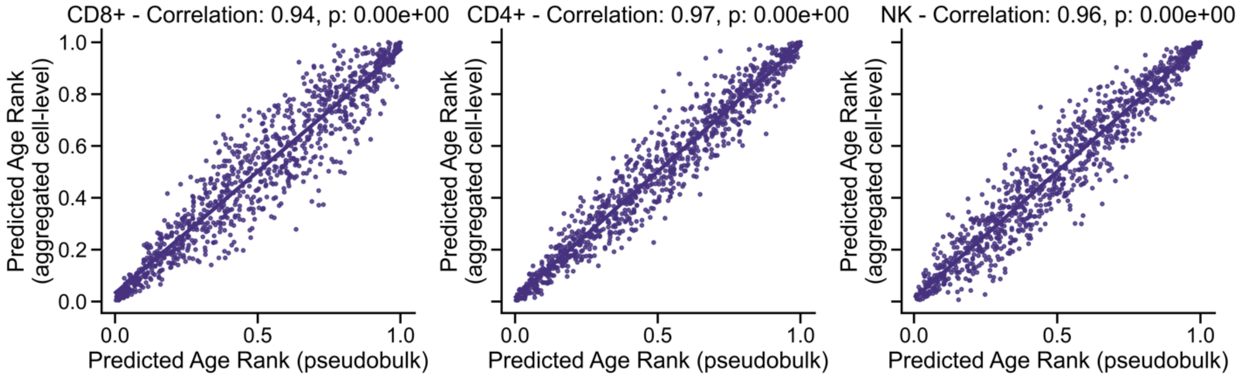


**Supplementary Figure S1. Validation of IMMClock's performance and reliability at single-cell resolution.** The figure shows the correlation between immune ages obtained by directly applying IMMClock to pseudobulk data (X-axis) and those inferred at single-cell resolution and subsequently aggregated to the patient level by averaging immune ages across all cells per patient (Y-axis). The subpanels display results for CD8+ cells, CD4+ cells, and NK cells (left to right), each using the respective mixed model. Analysis was conducted on the OneK1K cohort in cross-validation. Correlation values are 0.94 for CD8+ cells, 0.97 for CD4+ cells, and 0.96 for NK cells.

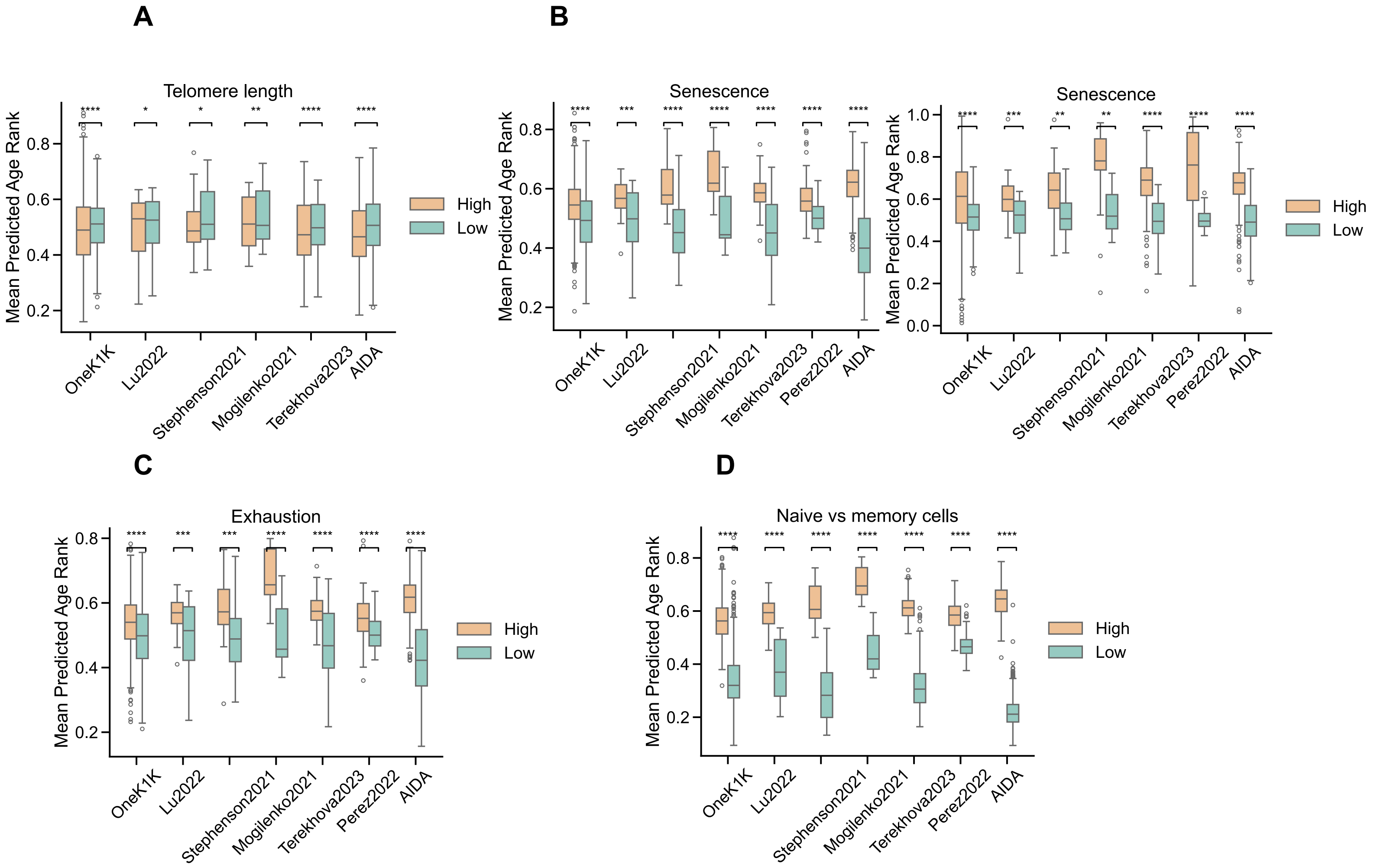


**Supplementary Figure S2. IMMClock associations with known $CD8^+$ T cell signatures at patient-level resolution.** This figure illustrates the association between predicted immune age and established T cell signatures of cellular phenotypes. We evaluated these associations using cross-validation on the OneK1K dataset and six independent scRNA-seq datasets: AIDA[28], Mogilenko et al. 2021[38], Terekhova et al. 2023[29], Stephenson et al, 2021[39], Perez et al. 2022[30], and Lu et al. 2022[26] The Y-axis represents the predicted immune age, calculated at the patient level by averaging immune age across all relevant cells per patient. Associations were tested using a paired Wilcoxon test, comparing the mean immune ages of high-expressing versus low-expressing cells across patients. **(A)** Higher immune age is associated with reduced expression of telomere-maintenance genes (TERT, DKC1). **(B)** Increased immune age corresponds to higher expression of senescence-related genes (KLRG1, B3GAT1). **(C)** Higher immune age is linked to increased expression of exhaustion markers (LAG3, TIGIT, PDCD1, HAVCR2, CTLA4). **(D)** IMMClock estimates that naïve cells have a lower immune age compared to memory cells. Statistical notation: **** $p < 0.0001$; *** $p < 0.001$; ** $p < 0.01$; * $p < 0.05$; ns, not significant.

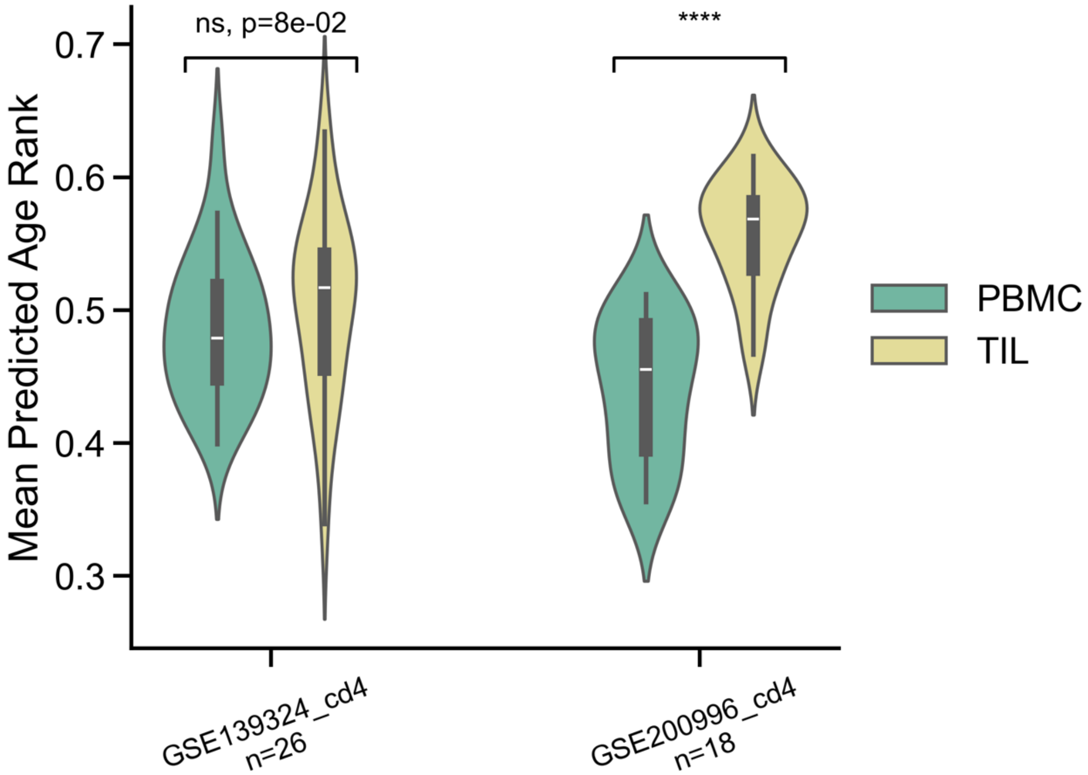


**Supplementary Figure S3.** IMMClock assessment of CD4⁺ immune age in tumor-infiltrating lymphocytes (TILs) compared to matched peripheral blood mononuclear cells (PBMCs) across two HNSCC cohorts (Ruffin et al., 2021; Luoma et al., 2022). The results suggest a trend toward higher predicted immune age for CD4⁺ T cells within the tumor microenvironment (TME) compared to blood samples, with statistical significance observed in one dataset (GSE200996, Ruffin et. al, p = 3.81e-06) but not the other (GSE139324, Luoma et al., p = 0.08). Statistical notation: **** p < 0.0001; *** p < 0.001; ** p < 0.01; * p < 0.05; ns, not significant.

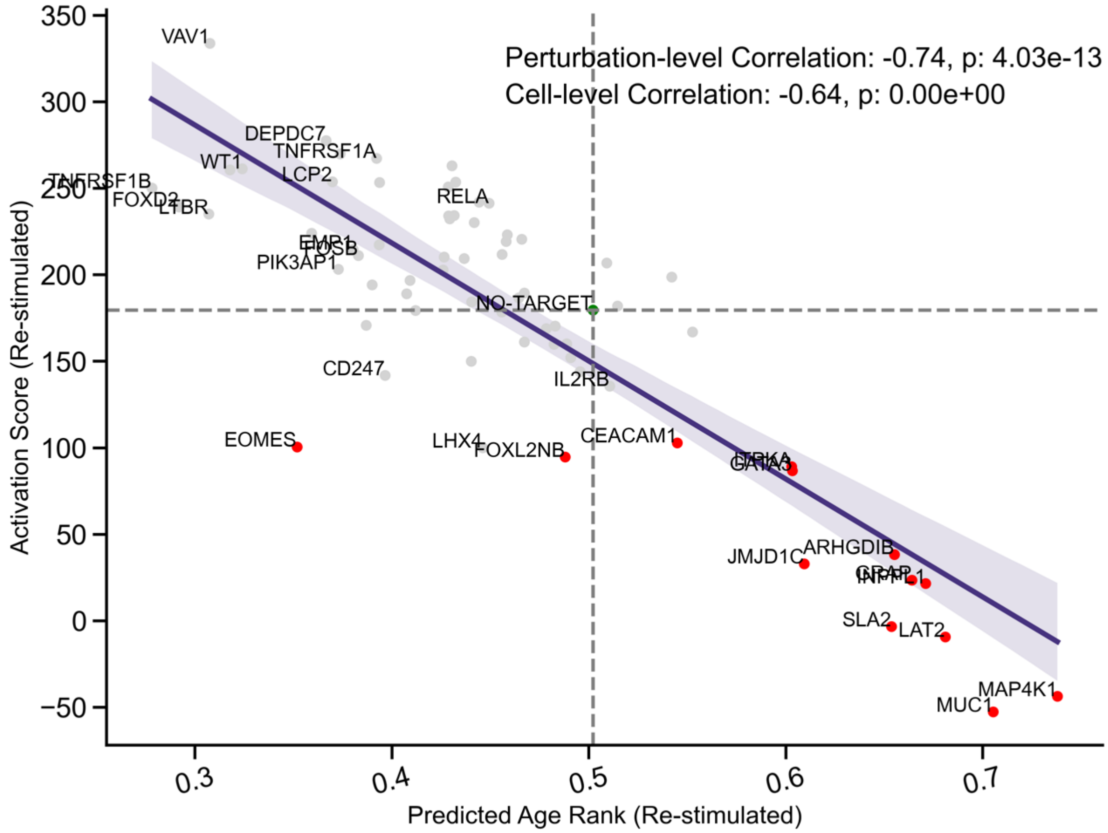


**Supplementary Figure S4. Younger T cells are associated with higher activation levels, even when controlling for cell cycle phase by focusing on G1 phase cells.** This figure, similar to Figure 6D in the main text, shows a strong negative correlation (ρ = -0.74) between immune age and activation score. Both the X-axis (mean predicted immune age) and Y-axis (mean activation score) represent aggregated values per perturbation across 70 gene perturbations**.**

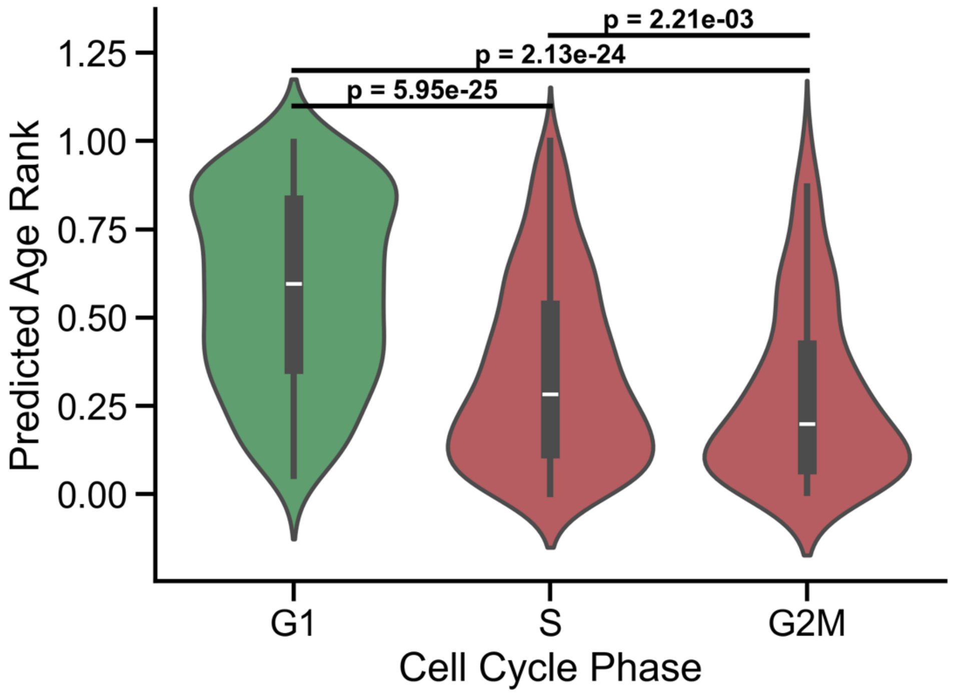


**Supplementary Figure S5.** The Y-axis represents the predicted immune age of individual NO-TARGET control $CD8^+$ T cells, displayed by cell cycle phase. $CD8^+$ T cells in cycling phases ($S/G_2M$) exhibit a younger immune age compared to those in the $G_1$ phase, indicating that cells actively progressing through the cell cycle tend to have a lower predicted immune age.
